# Hydroxide Ion Carrier for Proton Pump in Bacteriorhodopsin: Primary Proton Transfer

**DOI:** 10.1101/2019.12.23.887216

**Authors:** M. Imai, J. Ono, Y. Nishimura, H. Nakai

## Abstract

Bacteriorhodopsin (BR) is a model protein for light-driven proton pumps, where the vectorial active proton transport results in light-energy conversion. To clarify the microscopic mechanism of primary proton transfer from retinal Schiff base (SB) to Asp85 in BR, herein we performed quantum-mechanical metadynamics simulations of the whole BR system (∼3800 atoms). The simulations showed a novel proton transfer mechanism, viz. hydroxide ion mechanism, in which the deprotonation of specific internal water (Wat452) yields the protonation of Asp85 via Thr89, after which the resulting hydroxide ion accepts the remaining proton from retinal SB. Furthermore, systematic investigations adopting four sequential snapshots obtained by the time-resolved serial femtosecond crystallography revealed that proton transfer took 2–5.25 μs on the photocycle. The presence of Wat401, which is the main difference between snapshots at 2 and 5.25 μs, is found to be essential in assisting the primary proton transfer.

**SIGNIFICANCE:** Bacteriorhodopsin (BR), the benchmark of light-driven proton pumps, has attracted much attention from diverse areas in terms of energy conversion. Despite the significant experimental and theoretical efforts, the microscopic mechanism of the proton transfers in BR is not completely unveiled. In this study, quantum-mechanical molecular dynamics simulations of whole BR system were performed to elucidate the primary proton transfer in the L intermediate state with the latest snapshots obtained from X-ray free electron laser. As a result, it is found that the hydroxide ion originating from the specific internal water, which appears at the active site only in the L state, acts as a carrier for the primary proton transfer, demonstrating the importance of hydroxide ions in proton pumps.

## 1. INTRODUCTION

Proton transfer reactions play a vital role in various biological phenomena as well as chemical events. For example, the concentration gradient of protons across the membrane generated by the active transport through proton pumps becomes the driving force for adenosine triphosphate (ATP) synthesis, thus indicating that the vectorial proton transfers across the membrane help in energy conversion in biological systems.^1^ A great deal of effort has been devoted to elucidating the mechanism behind proton transfer reactions in biological and chemical macromolecules. However, the microscopic mechanism is still unclear due to the difficulties involved in direct observation and analysis of the proton transfer reactions, including the limitations imposed by high computational costs of quantum-mechanical simulations of biomolecules, and accurate measurements of the spatio-temporal resolution in experiments.

Bacteriorhodopsin (BR), shown in Figure 1, is a typical membrane protein that also functions as a proton pump.^1-3^ Recently, BR was adopted as a tool for efficient light-energy conversion in artificial cells.^4,5^ BR is considered a benchmark for microbial rhodopsins that have garnered attention from various research fields, owing to the diversity in functionality and applicability.^6-9^ For instance, part of microbial rhodopsins have been successfully utilized as molecular tools for controlling neuronal networks in optogenetics.^10,11^ In BR, starting from the photo-isomerization of the retinal chromophore^12-16^, which connects with Lys216 via the protonated Schiff base (SB), at least five proton transfer reactions successively take place on the photocycle, resulting in net proton translocation from the cytoplasmic to extracellular sides.^1-3,17-27^ In particular, the primary proton transfer occurring in the L intermediate state from the protonated SB to Asp85 is important, because it is the expected switching mechanism for preventing the backflow of protons.^22-25,27^

**Figure 1.**
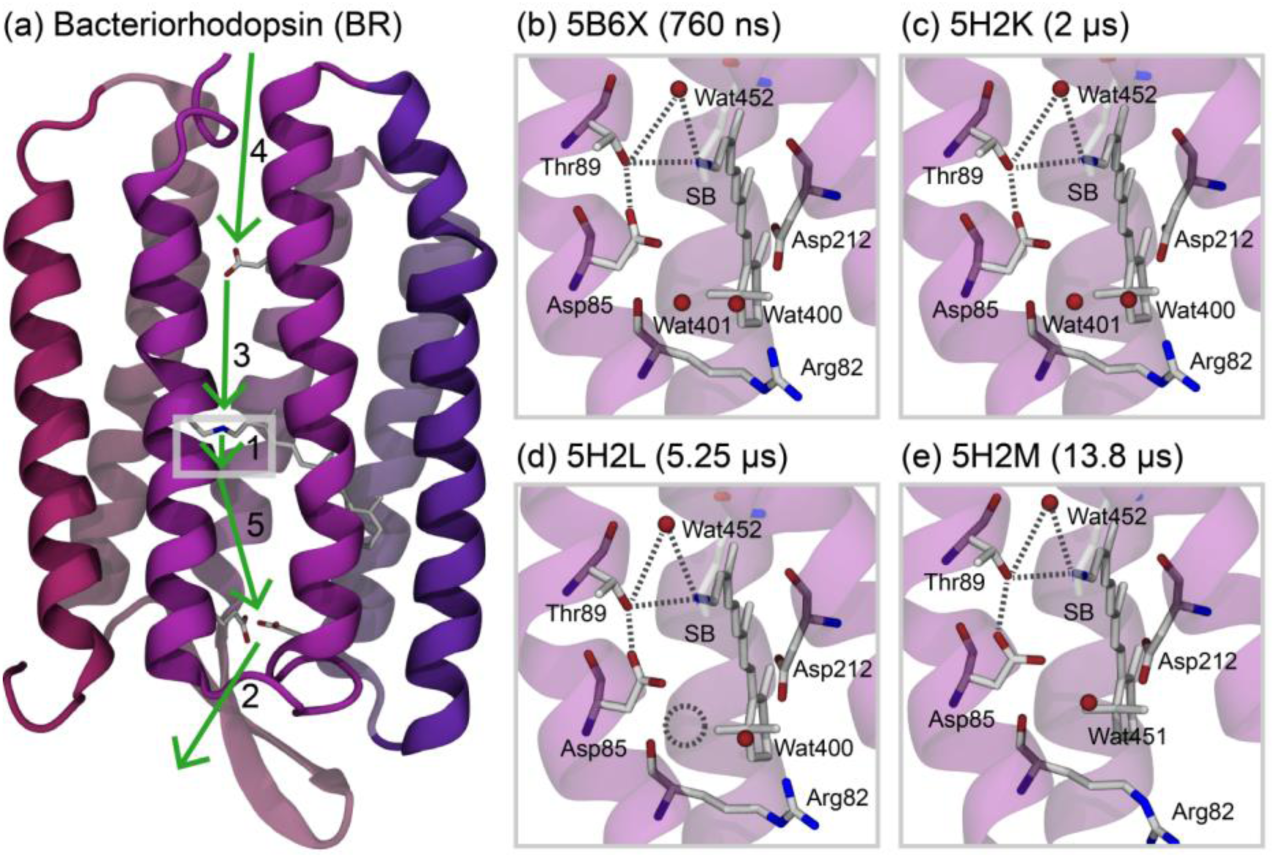
(a) Crystal structure of BR (PDB ID: 5H2K). The arrows indicate the consecutive proton transfer reactions in BR. The close-ups of the active site for the primary proton transfer in (b) 5B6X, (c) 5H2K, (d) 5H2L, and (e) 5H2M. The dotted lines indicate the putative hydrogen-bond network connecting the donor (SB) and acceptor (Asp85) for primary proton transfer. The dotted circle in 5H2L highlights the disappearance of the internal water (Wat401).

In 2016, E. Nango *et al*. reported a molecular movie of structural changes for BR along the photocycle captured by time-resolved serial femtosecond crystallography using an X-ray free electron laser (XFEL).^28^ This state-of-the-art technique is also utilized for capturing the molecular movie in the initial photoisomerization process of BR.^29,30^ In addition, time-resolved serial synchrotron crystallography has achieved the visualization of the slow structural changes for BR in the millisecond regime on the photocycle.^31^ These experimental frameworks induce the paradigm shift in structural biology from the conventional static pictures to the dynamical movies for (at least) photoreceptive proteins.^32^

The sequential 13 snapshots recorded by E. Nango *et al*. during the transition from the resting state to M state showed the functionally relevant conformational changes of BR, including rearrangements of internal water molecules in the active site and reorientational motions of the specific residues.^28^ The appearance of the internal water molecule (Wat452) exclusively in the L state (Figure 1b), the disappearance of the internal water molecule (Wat401) between 2 and 5.25 μs (Figure 1d), and the change in orientation of the Arg82 side chain between 5.25 and 13.8 μs (Figure 1e) are of special interest. However, the hydrogen atoms and protons are still invisible due to the restriction of spatial resolution (2.1 Å).^28^ Therefore, complementary approaches are expected to reveal the remaining mysteries in the reaction mechanism of proton transfers.

To this effect, a theoretical investigation based on quantum-mechanical molecular dynamics (QM-MD) simulations is one of the more convenient approaches. However, its application has been hampered by the so-called exponential wall problem in QM-MD computations with respect to the system size. Various efforts have been made to solve this problem.^33-35^ Although one of the feasible approaches might involve the first-principle simulations with pseudo potential for core electrons and plane-wave basis for valence electrons, the number of atoms that can be treated might be fewer than 1000 for pico or sub-pico second simulations.^36^

Recently, the authors’ group enabled to perform QM-MD simulations for thousands or tens of thousands of atoms by utilizing three pillars: linear-scaling divide-and-conquer (DC) technique, cost-effective semi-empirical density-functional tight-binding (DFTB) method, and massively parallel implementation with the hybrid of MPI and OpenMP.^37^ The DC-DFTB-MD simulations were successful in describing various chemical/physical phenomena in complex systems.^38-48^ The program system, DCDFTBMD^49^, has been distributed to the academic community through a special web site.^50^

In this study, we applied the DC-DFTB-MD method to examine the primary proton transfer in BR, in which the proton is transferred from the protonated SB to Asp85 during the L-to-M transition. This is the key step in achieving unidirectional active proton translocation. To elucidate the detailed pathway, timing, and the roles of residues and internal water molecules in the vicinity of the active site on the primary proton transfer in BR, we performed DC-DFTB-MD and metadynamics (metaD)^51-54^ simulations starting from four snapshots along the L-to-M transition in the molecular movie^28^, which are free from the radiation damages induced by x-ray, followed by analysis of the free energy surfaces.

## 2. METHODS

For crystal structures of BR in the L intermediate state on the photocycle, the four sequential snapshots obtained from time-resolved serial femtosecond crystallography based on XFEL were employed; the PDB IDs (times from photo-illumination) are 5B6X (760 ns), 5H2K (2 μs), 5H2L (5.25 μs), and 5H2M (13.8 μs), respectively.^28^ From these four snapshots, BR (the residues from 4 to 231), retinal, and internal water molecules given in the individual crystal structures (the B conformations) were extracted for constructing the isolated models. The missing internal coordinates in the N terminus (GLN1, ALA2, and GLN3) were added using the Swiss PDB Viewer.^55^ The protonation states of the titratable amino acids were evaluated at pH 7 with PROPKA^56^ implemented in the PDB2PQR package^57,58^, and as a result, Asp96, Asp115, and Glu194 were assigned to be protonated in these four structures, which is consistent with the previous Fourier transform infrared spectroscopy (FTIR) experiments^1^. In addition, the retinal SB was considered to be protonated in these four structures as the initial states of the primary proton transfer from SB to Asp85. The hydrogen atoms were added by the LEaP module in the Amber14 software.^59^ The resulting number of atoms for the isolated models (i.e., BR in vacuum) based on 5B6X, 5H2K, 5H2L, and 5H2M were 3748, 3748, 3745, 3742, respectively.

The positions of the complemented missing residues (the residues from 1 to 3) and the hydrogen atoms in the systems were first relaxed by energy minimization with the restraint of the non-hydrogen atoms in the remaining residues within the framework of the classical force fields. The minimization process was conducted using the Sander module of Amber 14^59^, with the Amber ff99SB-ILDN force field^60^ for the standard amino acids, and TIP3P^61^ for the internal water molecules. The parameters for the 13-*cis* retinal bound to Lys216 through SB were generated with the RESP charges at the B3LYP/6-31G(d) level and the general Amber force field (GAFF).^62^ The resulting relaxed structures were used as the initial structures for the DC-DFTB-MD simulations, in which all the atoms (approximately 3,800 atoms) were treated quantum mechanically without any restraints.

The DC-DFTB-MD simulations were performed for the equilibration and production runs of the systems using the DCDFTBMD program.^49^ The DFTB3 method was employed with the 3OB parameter set.^63-65^ Grimme’s DFT-D3 dispersion correction with the Becke-Johnson damping scheme was adopted.^66,67^ The DC method with 4 × 4 × 4 Å^3^ cubic grids, resulting in ca. 660 subsystems, and the buffer radius of 5 Å was employed to reduce the computational costs. The unbiased DC-DFTB-MD simulations starting from the above-mentioned structures were carried out for 50 ps under the *NVT* ensemble (*T* = 298.15 K) controlled by the Andersen thermostat.^68^ Here, the velocity Verlet integrator with a time step of 0.5 fs was utilized for the time integration. From the 50-ps DC-DFTB-MD simulations for equilibrating the systems, no proton transfers were actually observed. The structural analyses such as the probability distribution functions for the hydrogen-bond distances and angles in the active site were performed on the basis of the last 10-ps trajectories.

The free energy surfaces (FESs) for the primary proton transfer in BR were evaluated using the DC-DFTB-metaD simulations.^49^ In the present study, the well-tempered metadynamics (WTmetaD)^52^ was employed to accelerate the deprotonation event of the protonated SB. As the collective variable (CV) of one-dimensional (1D) WTmetaD characterizing the dissociation event of the protonated SB, the effective coordination number for the N atom on SB with respect to proton was selected, which is defined as the rational form as a function of the distance between N and H atoms of SB with the cutoff distance of 1.6 Å.^49^ The initial height of the Gaussian function for the biasing potential was set to 1.88 kcal/mol. Note that this value of the initial Gaussian height was actually employed in the previous quantum-mechanics/molecular-mechanics (QM/MM) metaD simulations for biomolecular systems.^69^ The width of the Gaussian function was set to 0.02. This value corresponds to the standard deviation of CV obtained from unbiased MD simulations in the initial state of the primary proton transfer.^53^ The Gaussian biasing potential was deposited at the time interval of 20 fs, which is estimated from the initial decay of the autocorrelation function for CV in the unbiased MD simulations.^53^ The bias factor of WTmetaD was set to 31 corresponding to the bias temperature of 9000 K.

The two-dimensional (2D) WTmetaD were performed to confirm whether the protonation change of the internal water molecule in the cytoplasmic side of the active site (Wat452) occurs before the deprotonation of SB. In addition to the coordination number for the N atom on SB employed in the 1D WTmetaD, that for the O atom on Wat452 with the cutoff distance of 1.4 Å was adopted as the alternate CV in the 2D WTmetaD. The parameters for the 2D WTmetaD were identical with those for the 1D WTmetaD.

To characterize the proton transfer reaction unambiguously, 2D FESs for the charge fluctuations of the donor (SB) and acceptor (Asp85), instead of the corresponding coordination numbers, were calculated. Note that recently the charge fluctuation is proposed as the efficient CV for the FESs of chemical reactions by performing the coupled-perturbed DFTB-metaD simulation.^70^ In this study, the reweighting method for WTmetaD^71^ was employed to obtain the 2D FESs for the charge fluctuations from the DC-DFTB-metaD with the CV(s) of the coordination number(s). Here, the charge fluctuation of SB *q*_SB_ is defined as

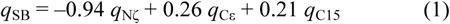

where *q*_Nζ_, *q*_Cε_, and *q*_C15_ denote the charge of the N atom on SB, that of the Cε atom in Lys216, and that of the C15 atom in retinal, respectively. Similarly, the charge fluctuation of Asp85 *q*_D85_ is defined as

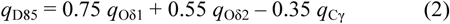

where *q*_Oδ1_, *q*_Oδ2_, and *q*_Cγ_ indicate the charge of the Oδ atom in Asp85 forming hydrogen bond with Thr89, that of the remaining Oδ atom in Asp85, and the Cγ atom in Asp85, respectively. Note that these coefficients were determined from the principal component analysis for the time series of the charges. The weighted 2D histograms for *q*_SB_ and *q*_D85_ were calculated from multiple biased trajectories using the reweighting method,^71^ and the resulting histograms were statistically averaged with weighted histogram analysis method (WHAM).^72^ After equilibration with DC-DFTB-MD, at least 5 biased trajectories with different initial structures and velocities were generated from DC-DFTB-WTmetaD for more than 50 ps during which the 2D FES for the charge fluctuations *q*_SB_ and *q*_D85_ were calculated.

## 3. RESULTS AND DISCUSSION

First, we performed the DC-DFTB-metaD simulations biasing the deprotonation of SB: i.e., 1D metaD with a single CV. This is based on the assumption that deprotonation initiates the primary proton transfer as examined in the previous studies.^22-25^ We then obtained 17 proton transfer trajectories from SB to Asp85 by using the 5H2K structure, which were categorized into two parts. One is a two-step proton transfer pathway, denoted as path **1**, in which the proton migration occurs from SB to the hydroxy group in the side chain of Thr89, followed by the proton transfer from Thr89 to Asp85 (Figure 2a). Path **1** was observed in 8 out of 17 metaD trajectories. This pathway was proposed by the NMR study.^27^ The other is a three-step proton transfer pathway, denoted as path **2**, where the protonation of Wat452 is first induced by the deprotonation of SB, and then the proton is transferred from Wat452 through Thr89 to Asp85 (Figure 2b). Here, Wat452 plays a key role in the initial proton transfer by producing hydronium ion (H_3_O^+^), followed by deprotonation, which corresponds to the standard Grotthuss hopping.^73^ Path **2** was observed in 9 out of 17 metaD trajectories. This pathway was proposed by the XFEL study.^28^ The structural condition that brought about path **2** might be the formation of simultaneous short-range hydrogen bonds from SB to Asp85 through Wat452. The detailed analysis is given in Figures S1 and S2 in the supporting information (SI).

**Figure 2.**
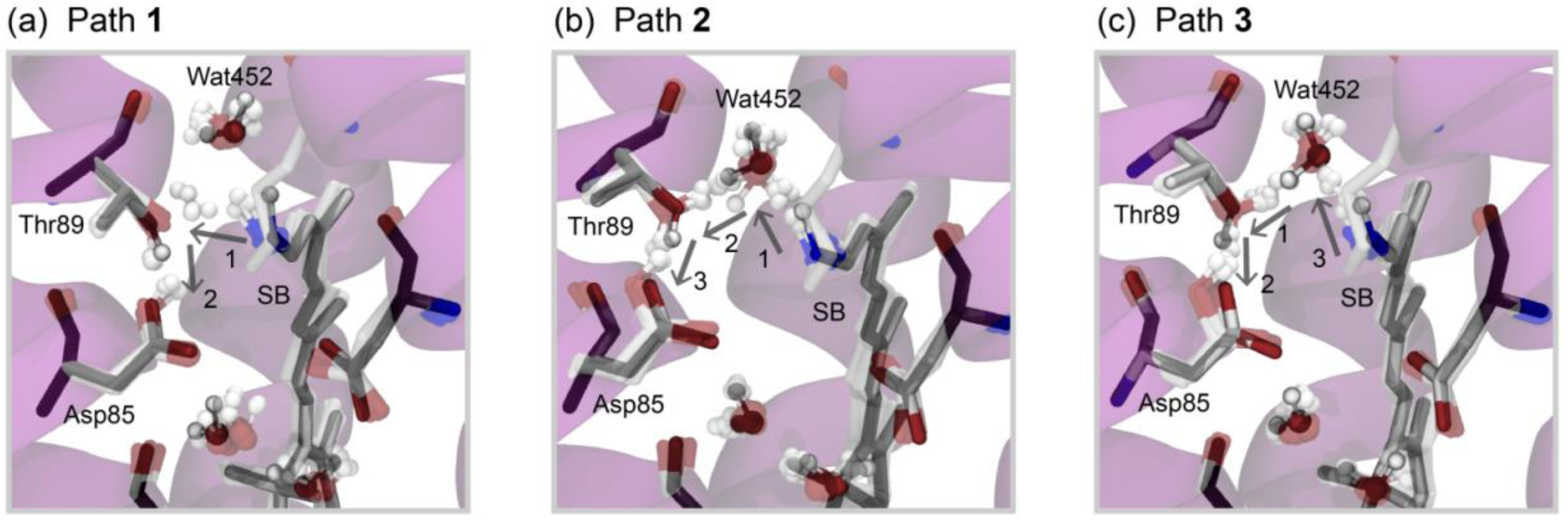
Representative snapshots of the primary proton transfer process in BR obtained from DC-DFTB-metaD simulations with the 5H2K structure for (a) path **1**, (b) path **2**, and (c) path **3**. Here, successive ten structures along the metaD trajectories with the intervals of 100 fs were superimposed. The arrows and numbers indicate the direction and order of the proton transfers, respectively.

Figures 3a and 3b show the 2D FESs for two paths **1** and **2**, both of which were obtained from the DC-DFTB-metaD simulations using the 5H2K structure. Here, the horizontal and vertical axes represent the charge fluctuations of SB and Asp85, *q*_SB_ and *q*_D85_, respectively. These values are recognized as the order parameters of the protonation states for SB and Asp85. The free energy minimum located at {*q*_SB_, *q*_D85_} ≃ {0.1, −1.3} corresponds to the initial state (i.e., protonated SB and deprotonated Asp85), whereas the one located at {*q*_SB_, *q*_D85_} ≃ {0.4, −1.0} corresponds to the final state (i.e., deprotonated SB and protonated Asp85). The intermediate state (i.e., deprotonated SB and deprotonated Asp85) is located at around {0.4, −1.3}, and represents protonated Thr89 in paths **1** and **2**, as well as the hydronium ion of Wat452 in path **2**. The free energy barriers from the initial to final states, Δ*G*^‡^, of paths **1** and **2** were estimated to be 59 and 58 kcal/mol, respectively. From the timescale of the L state, i.e., the order of microseconds,^1^ the barrier can be estimated to be approximately 10 kcal/mol using the transition state theory. Therefore, the present estimations of Δ*G*^‡^ of paths **1** and **2** are significantly higher than the previous theoretical values.^22-25^

**Figure 3.**
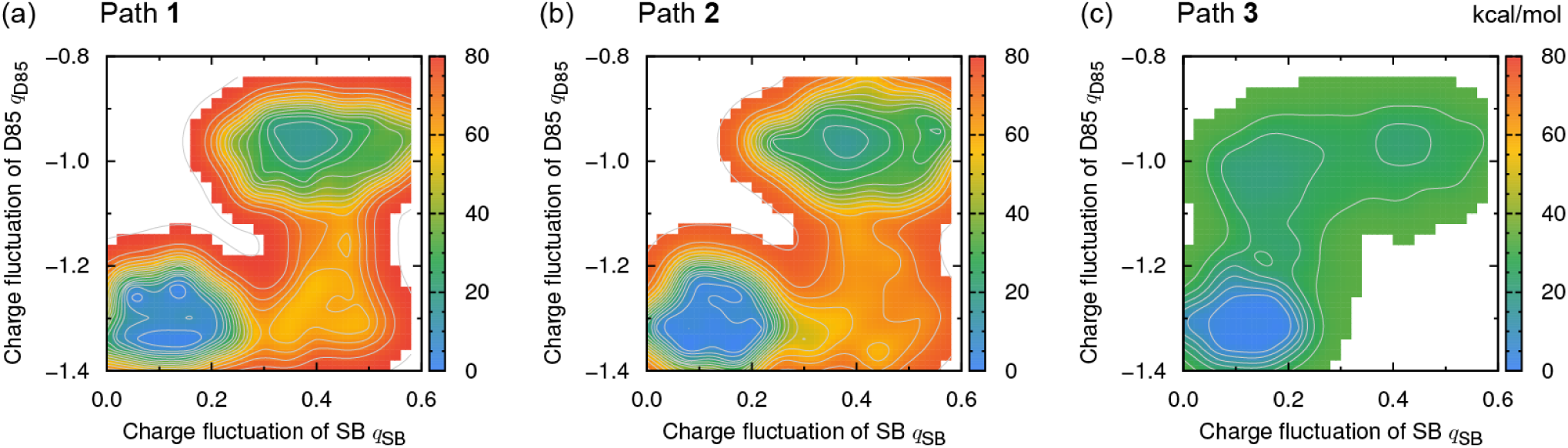
Two-dimensional free energy surfaces in kcal/mol projected to the charge fluctuations of SB *q*_SB_ and Asp85 (D85) *q*_D85_ obtained from DC-DFTB-metaD simulations and the reweighting method for the 5H2K structure of BR: (a) path **1**, (b) path **2**, and (c) path **3**.

Next, we performed the DC-DFTB-metaD simulations with the biases for coordination numbers of SB and Wat452; i.e., 2D metaD with two CVs. It is notable that the simulations did not assume that the deprotonation of SB precedes the other protonation changes in the active site. From the simulations with the 5B6X and 5H2K structures, we obtained another three-step proton transfer pathway, denoted as path **3**, in which the deprotonation of Wat452 gives rise to the protonation of Asp85 via Thr89, followed by the reprotonation of the resultant hydroxide ion of Wat452 from SB (Figure 2c). Because the hydroxide ion (OH^-^) of Wat452 plays a key role in this pathway, we termed this as the hydroxide ion mechanism. Note that in contrast to the above-mentioned paths **1** and **2**, the hydroxide ion mechanism via path **3** had not been discovered so far, to the best of our knowledge. The structural analysis in path **3**, as shown in Figure S3 in the SI, clarified that short-range hydrogen bonds related with Wat452 switches stepwise from Thr89-water to SB-hydroxide ion, which might be caused by the higher mobility of Wat452. We speculate that this stepwise switching of the hydrogen bond might be the origin of the irreversible proton transfer.

Figure 3c shows the 2D FES for path **3** obtained from the DC-DFTB-metaD simulations with the 5H2K structure. The free energy minima corresponding to the initial and final states in path **3** are the same as those in paths **1** and **2**. The local minimum located at {*q*_SB_, *q*_D85_} ≃ {0.1, −1.0} indicates the intermediate state (i.e., protonated SB and protonated Asp85) with the hydroxide ion of Wat452. The free energy barrier from the initial to final states Δ*G*^‡^ in path **3** was estimated to be 29 kcal/mol, which was slightly higher than those in the previous QM/MM studies^22-25^ based on SCC-DFTB with crystal structures including x-ray-induced radiation damages^28^, but significantly lower than those in paths **1** and **2**. Therefore, the present simulations indicate that the hydroxide ion mechanism is the most favorable from an energetic standpoint.

Table 1 summarizes the free energy barriers Δ*G*^‡^ of the primary proton transfer in paths **1, 2**, and **3** for 5B6X, 5H2K, 5H2L, and 5H2M. Note that in 5H2L and 5H2M the proton transfer via path **3** was not obtained from the 2D metaD simulations. Instead, the irrelevant proton transfer from Wat452 to the backbone carbonyl group of Thr89 was observed, as shown in Figure S4. The value of Δ*G*^‡^ in path **3** for 5H2K was the lowest among the three pathways and four L-state structures, indicating that the primary proton transfer takes place from 5H2K (2 μs) to 5H2L (5.25 μs) on the photoreaction cycle in BR.

**Table 1.** Free energy barriers Δ*G*^‡^ in kcal/mol obtained from the 2D FESs for each pathway of the primary proton transfer in BR.

|  | 5B6X | 5H2K | 5H2L | 5H2M |
| --- | --- | --- | --- | --- |
| Path 1 | 63 | 59 | 60 | 56 |
| Path 2 | 62 | 58 | 64 | 64 |
| Path 3 | 30 | 29 | - | - |

Finally, we examined the differences in the four L-states (5B6X, 5H2K, 5H2L, and 5H2M) from a structural viewpoint. In the hydroxide ion mechanism via path **3**, the hydrogen bond between Wat452 and the side-chain hydroxy group of Thr89 should be essential for the initial deprotonation of Wat452. Figures 4a-d show the 2D probability distribution functions for the hydrogen-bond distance, *r*(O…O), and the hydrogen-bond angle, *θ*(O-H…O), which were evaluated by using the results of DC-DFTB-MD simulations with the 5B6X, 5H2K, 5H2L, and 5H2M structures at the equilibrium condition. In the 5B6X and 5H2K structures the peaks of the probability distributions were located at {*r, θ*} = {2.7 Å, 160°}, whereas in the 5H2L and 5H2M structures the peaks were at {3.9 Å, 90°}. These results indicate that the hydrogen bond between Wat452 and Thr89 in 5B6X/5H2K was favorably formed, while that in 5H2L/5H2M was broken; in the latter case, Wat452 formed the hydrogen bond with the main-chain carbonyl group of Thr89. Note that there are no significant differences in the other hydrogen bonds connecting SB and Asp85 in the active site amongst the four structures under discussion (Figures S5, S6, and S7).

**Figure 4.**
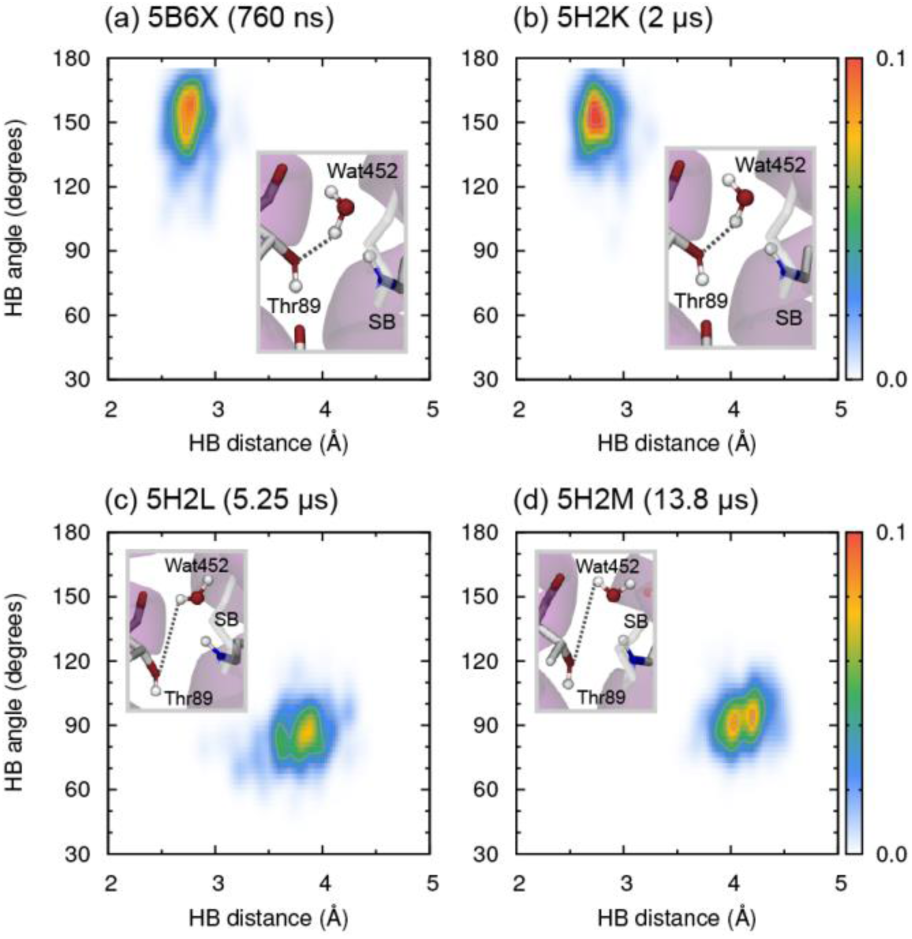
Two-dimensional probability distribution functions for the hydrogen-bond (HB) distance and the HB angle between Wat452 and Thr89.

One of the prominent differences between 5B6X/5H2K and 5H2L/5H2M is the number of internal water molecules in the extracellular side of the active site: two and one, respectively (Figure 1). To clarify the role of Wat401, we performed the DC-DFTB-MD simulation with the 5H2K structure by artificially eliminating Wat401. The 2D probability distribution function for the hydrogen-bond distance and angle obtained from the simulation closely resembled that in 5H2L/5H2M (Figure S8). Therefore, Wat401 was found to assist the initial proton transfer in path **3** by stabilizing the hydrogen bond between Wat452 and Thr89.

## 4. CONCLUSION

The present study performed the full QM-MD/metaD simulations for the primary proton transfer in BR using the DC-DFTB method, in which the whole system was treated quantum-mechanically. From the 1D metaD simulations based on the conventional assumption that the deprotonation of SB occurs prior to the other protonation changes, two pathways (paths **1** and **2**) were observed, as predicted in the previous studies.^22-25,27,28^ However, free energy barriers of both pathways were estimated to be significantly higher. In contrast, from the 2D metaD simulations biasing the protonation changes of both SB and Wat452, a hitherto unknown pathway (path **3**) was observed. In this pathway, Wat452 initially donates the proton to Asp85 via Thr89, and subsequently the resulting hydroxide ion receives the proton from SB. Here, Wat452 was expected to serve as the switch from Thr89 to SB of the short-range hydrogen bonds required in the transition states of the proton transfers, which could be biologically relevant to the unidirectional proton transfers as the proton pump. From energetic and structural viewpoints, the present results confirmed that the primary proton transfer occurs from 5H2K (2 μs) to 5H2L (5.25 μs) via path **3**, which we call the hydroxide ion mechanism. After the primary proton transfer, Wat401 disappears from the extracellular side of the active site, which destabilizes the hydrogen bond between Wat452 and Thr89 and further discourages the reverse reaction. Finally, the downward orientation of Arg82 promotes the next proton transfer process, i.e., the release of the proton into the extracellular medium^73-78^, and the subsequent reprotonation of SB from Asp96 on the cytoplasmic side.^79-81^

## SUPPORTING MATERIAL

Supporting materials can be found online at https://doi.org/10.1016/j.bpj.***.

## AUTHOR CONTRIBUTIONS

M.I., J.O., and H.N. designed the research. Y.N. developed the simulation tools. J.O. developed the analytical tools. M.I. and J.O. performed the calculations and analyses. J.O. and H.N. wrote the paper. H.N. supervised the entire project. All authors reviewed the results and approved the final version of the manuscript.

## ACKNOWLEDGMENTS

This work was supported in part by a Grant-in-Aid for Scientific Research (A) “KAKENHI Grant Number JP26248009” and a Grant-in-Aid for Scientific Research (S) “KAKENHI Grant Number JP18H05264” from the Japan Society for the Promotion of Science (JSPS). The calculations were in part performed at the K computer provided by the RIKEN Advanced Institute for Computational Science through the HPCI System Research project (Project ID: hp170040) and at Research Center for Computational Science (RCCS), Okazaki Research Facilities, National Institutes of Natural Sciences (NIIS), Japan.

